# Oncogenic role of a developmentally regulated *NTRK2* splice variant

**DOI:** 10.1101/2022.01.07.475392

**Authors:** Siobhan S. Pattwell, Sonali Arora, Nicholas Nuechterlein, Michael Zager, Keith R. Loeb, Patrick J. Cimino, Nikolas C. Holland, Noemi Reche-Ley, Hamid Bolouri, Damian A. Almiron Bonnin, Frank Szulzewsky, Vaishnavi V. Phadnis, Tatsuya Ozawa, Michael J. Wagner, Michael C. Haffner, Junyue Cao, Jay Shendure, Eric C. Holland

## Abstract

Temporally-regulated alternative splicing choices are vital for proper development yet the wrong splice choice may be detrimental. Here we highlight a novel role for the neurotrophin receptor splice variant TrkB.T1 in neurodevelopment, embryogenesis, transformation, and oncogenesis across multiple tumor types in both humans and mice. TrkB.T1 is the predominant *NTRK2* isoform across embryonic organogenesis and forced over-expression of this embryonic pattern causes multiple solid and nonsolid tumors in mice in the context of tumor suppressor loss. TrkB.T1 also emerges the predominant *NTRK* isoform expressed in a wide range of adult and pediatric tumors, including those harboring TRK fusions. Affinity purification-mass spectrometry (AP-MS) proteomic analysis reveals TrkB.T1 has distinct interactors with known developmental and oncogenic signaling pathways such as Wnt, TGF-ß, Hedgehog, and Ras. From alterations in splicing factors to changes in gene expression, the discovery of isoform specific oncogenes with embryonic ancestry has the potential to shape the way we think about developmental systems and oncology.

## Main

Embryogenesis, and neurodevelopment in particular, comprise an elegant and well-orchestrated series of tightly regulated events, culminating in an organized and highly functioning organism. Cancer, on the other hand, can often be viewed as an uncontrolled, unrestrained, genetically chaotic disease, lacking the precise spatial and temporal rigidity associated with normal development. While appearing to be on opposite sides of the organizational spectrum, the similarities between early development and oncogenesis are numerous. This has been observed frequently as key developmental signaling pathways, such as Wnt, Hedgehog, or Notch, have been shown to be dysregulated in cancer across all stages from tumor initiation and maintenance to metastasis (Aiello and Stanger 2016).

Encoded by the *NTRK2* gene, the tropomyosin receptor kinase TrkB, which has many alternatively spliced isoforms, has well-established roles across neurodevelopment, astrocyte biology, and has also been implicated in a wide range of cancer types (Nakagawara et al. 1994; Eggert et al. 2001; Pearse et al. 2005; Sclabas et al. 2005; Yu et al. 2008; Li et al. 2009; Kupferman et al. 2010; Lai et al. 2010; de Farias et al. 2012; Lee et al. 2012; Makino et al. 2012; Okamura et al. 2012; Bao et al. 2013; Sinkevicius et al. 2014; Kim et al. 2015; Vaishnavi et al. 2015; Yin et al. 2015; Akil et al. 2016; Yuzugullu et al. 2016; Zhang et al. 2017; Gomez et al. 2018; Moriwaki et al. 2018). *NTRK2* gene fusions have recently been the focus of many clinical and pharmacological studies (Cocco et al. 2018; Albert et al. 2019; Gatalica et al. 2019; Solomon and Hechtman 2019) while inhibition of the TrkB kinase has been shown to inhibit cell proliferation and contribute to apoptosis (Makino et al. 2012). In addition to well known-roles in neurobiology, recent studies suggest that *NTRK2* may render cells resistant to anoikis and prone to metastasis while others suggest its involvement in the epithelial-to-mesenchymal transition associated with increased migration and invasiveness of many cancer cell lines (Douma et al. 2004; Geiger and Peeper 2005; Sclabas et al. 2005; Desmet and Peeper 2006; Kupferman et al. 2010; Smit and Peeper 2011; Bao et al. 2013; Contreras-Zarate et al. 2019), yet the precise mechanisms of how TrkB exerts its oncogenic role are not fully known. While prior studies have been fundamental in uncovering *NTRK2* involvement in cancer, the complex post-translational modifications, intricate splicing patterns, and prior roles in embryogenesis are often ignored.

Recent findings have shown that a kinase-deficient *NTRK2* splice variant, TrkB.T1, is the predominant neurotrophin receptor in human gliomas, enhances glioma aggressiveness in mice, and increases the perdurance of PDGF-induced Akt and STAT3 signaling (Pattwell et al. 2020a). This splice variant contains a unique terminal exon encoding a 11-amino acid tail that is 100% conserved across species including chicks, mice, rats, felines, and humans (Middlemas et al. 1991; Klein et al. 1993; Biffo et al. 1995; Shelton et al. 1995; Baxter et al. 1997; Luberg et al. 2010), suggesting a potential evolutionarily significant biological role. In light of recent findings implicating this kinase-deficient splice variant in glioma biology (Pattwell et al. 2020a), we explore the role of TrkB.T1 in neural development and early embryonic central nervous system (CNS) and mesenchymal development in particular.

In this study, we characterize the role of the *NTRK2* splice variant, TrkB.T1, in embryonic neurodevelopment and early organogenesis via transcript analysis of an enhanced single-cell combinatorial-indexing RNA-sequencing analysis method (sci-RNA-seq3) data of E9.5 to E13.5 mouse embryos (Cao et al. 2019) and transcript specific immunohistochemistry. We demonstrate a developmental role for TrkB.T1 beyond that of the central nervous system, highlight TrkB.T1’s transformation potential *in vitro*, and uncover its role as an oncogenic driver *in vivo* using a RCAS/tv-a mouse model (Holland and Varmus 1998; Ozawa et al. 2014). We further reveal TrkB.T1 to be the predominant TRK isoform expressed across the majority of adult and pediatric tumors using clinical data from The Cancer Genome Atlas (TCGA) (Cancer Genome Atlas Research et al. 2013) and the Therapeutically Applicable Research to Generate Effective Treatments (TARGET) initiative, respectively, and uncover a several potential interactors via proteomic analysis. Using TrkB.T1, we explore the role of developmentally regulated splicing choices and subsequently show that forced expression of a specific isoform can be neoplastic in the same postnatal organs that once expressed it during embryogenesis.

## Results

### TrkB.T1 in embryonic development and organogenesis

It is widely known that the majority of organ development in rodents occurs prior to E18 (Cao et al. 2019) and recent advances in next generation sequencing have allowed for the characterization of transcriptional dynamics across development at a single cell resolution resulting in a Mouse Organogenesis Cell Atlas (MOCA) (Cao et al. 2019). To explore the expression of TrkB.T1 in CNS development, we utilized the fact that the TrkB.T1 and full-length TrkB.FL *NTRK2* transcripts differ in their 3’ ends(Luberg et al. 2010), which allows for transcript quantification in single cell RNA sequencing data and mapped expression of individual transcripts in mouse embryos over developmental time. Using sci-RNA-seq3 data, the MOCA (Cao et al. 2019) provides gene expression data and cell trajectory annotation for 2,026,641 cells from 61 mouse embryos across five development stages (E9.5-E.13.5), resulting in 38 main cell clusters and 10 main trajectories. The expression of transcripts for all protein coding genes, including TrkB.T1 and TrkB.FL, were normalized and visualized over the existing landscape provided by Cao et al (Cao et al. 2019). First, we examined TrkB.FL expression patterns in the *t*-SNE plot of all 38 cell types within the MOCA and found that TrkB.FL expression is appreciably restricted to a subset of cell types, all of which were neuronal in nature (**Fig. 1A, 1B, Supplemental Fig. S1A – S1E**). TrkB.T1, however, was expressed broadly across many different cell types within the embryo during this period (**Fig. 1C, 1D, Supplemental Fig. S1F – S1J**). We next sought to explore the *NTRK2* transcript expression patters within the CNS developmental trajectory and found that TrkB.T1 cellular expression is observed across multiple neurodevelopmentally driven cell clusters including radial glia, immature oligodendrocytes, isthmic organizer cells (**Fig. 1E**). The second largest population of cells with high, variable levels of TrkB.T1 expression during this developmental period is found in the mesenchymal trajectory, and mapping expression of both transcripts showed a vast predominance of TrkB.T1 over TrkB.FL similar to the non-neuronal cell types of the developing CNS (**Fig. 1E, 1F, 1G, 1H**). Visualization of *NTRK2* expression reveals intense, diffuse patterns of TrkB.T1 across multiple days in multiple cell types, with expression levels rivaling those of actin in particular clusters (**Supplemental Fig. S1F – S1J, Supplemental Fig. S2**), whereas TrkB.FL expression remains predominantly restricted to mature neurons (**Supplemental Fig. S1A – S1E**). An interactive site allowing 3D visualization and further exploration of NTRK2 transcripts among cell types and within the CNS and mesenchymal trajectories over developmental time can be found at https://atlas.fredhutch.org/fredhutch/ntrk2/.

**Figure 1:**
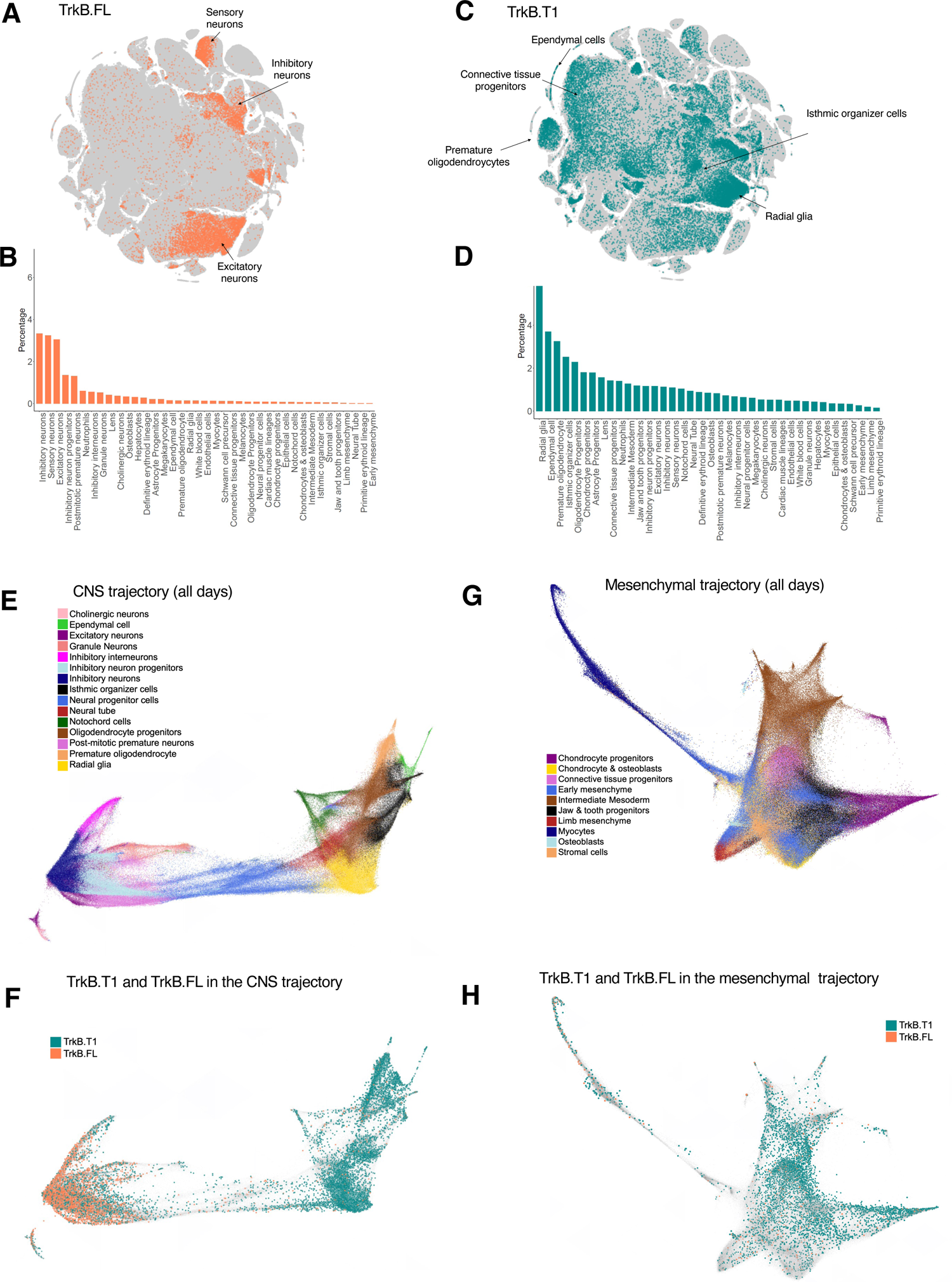
sci-RNA-Seq 3 transcript analysis across E9.5-E13.5 mouse embryonic development shows TrkB.FL expression in neuronal clusters with widespread TrkB.T1 expression in CNS and mesenchymal trajectories. *t*-SNE visualization of TrkB.FL expression (A,B) and TrkB.T1 (B,C) of 2,026,641 mouse embryo cells from E9.5-E13.5(Cao et al. 2019) shows high TrkB.FL expression in neuronal clusters and CNS clusters with strong, diffuse TrkB.T1 expression across the majority of cell clusters. Visualization of CNS (D) and mesenchymal (D) cell type trajectories across all days show strong expression of TrkB.FL in mature neuronal cells (F) with diffuse expression of TrkB.T1 in large subsets of immature stem-like cells across developmental trajectories (G).

As the sci-RNA-seq3 data show widespread TrkB.T1 expression across multiple clusters within and outside the CNS, we wanted to visualize TrkB.T1 and TrkB.FL expression across embryonic development beyond E13.5. To this end, we stained histological sections of embryos from E10-E17 with antibodies specific to the TrkB.FL kinase domain as well as an antibody specific to the intracellular 11-amino acid tail of TrkB.T1 (Pattwell et al. 2020b). Similar to the sci-RNA-seq3 data, examination at early embryonic time points, when the most rapid cellular growth and organogenesis is underway, the protein ratio between these two receptors is different, with TrkB.T1 present at exceedingly high levels in multiple organ sites (**Fig. 2A – 2I**), compared to TrkB.FL (**Fig. 2J – 2R**), which is predominantly expressed and maintained in CNS consistent with the transcript levels for the two TrkB gene products shown in **Fig. 1**. By the end of embryonic development circa E17, immunohistochemistry of both splice variants reveals that TrkB.T1 levels have decreased but are still present throughout various non-CNS organs, while the majority of TrkB.FL remains restricted within the CNS (**Fig. 2**). These data suggest that in addition to a role in CNS development and gliomagenesis, TrkB.T1 may have a more widespread role in early organogenesis, implying that it may also play a role in abnormal developmental patterns typically seen in non-CNS cancers.

**Figure 2:**
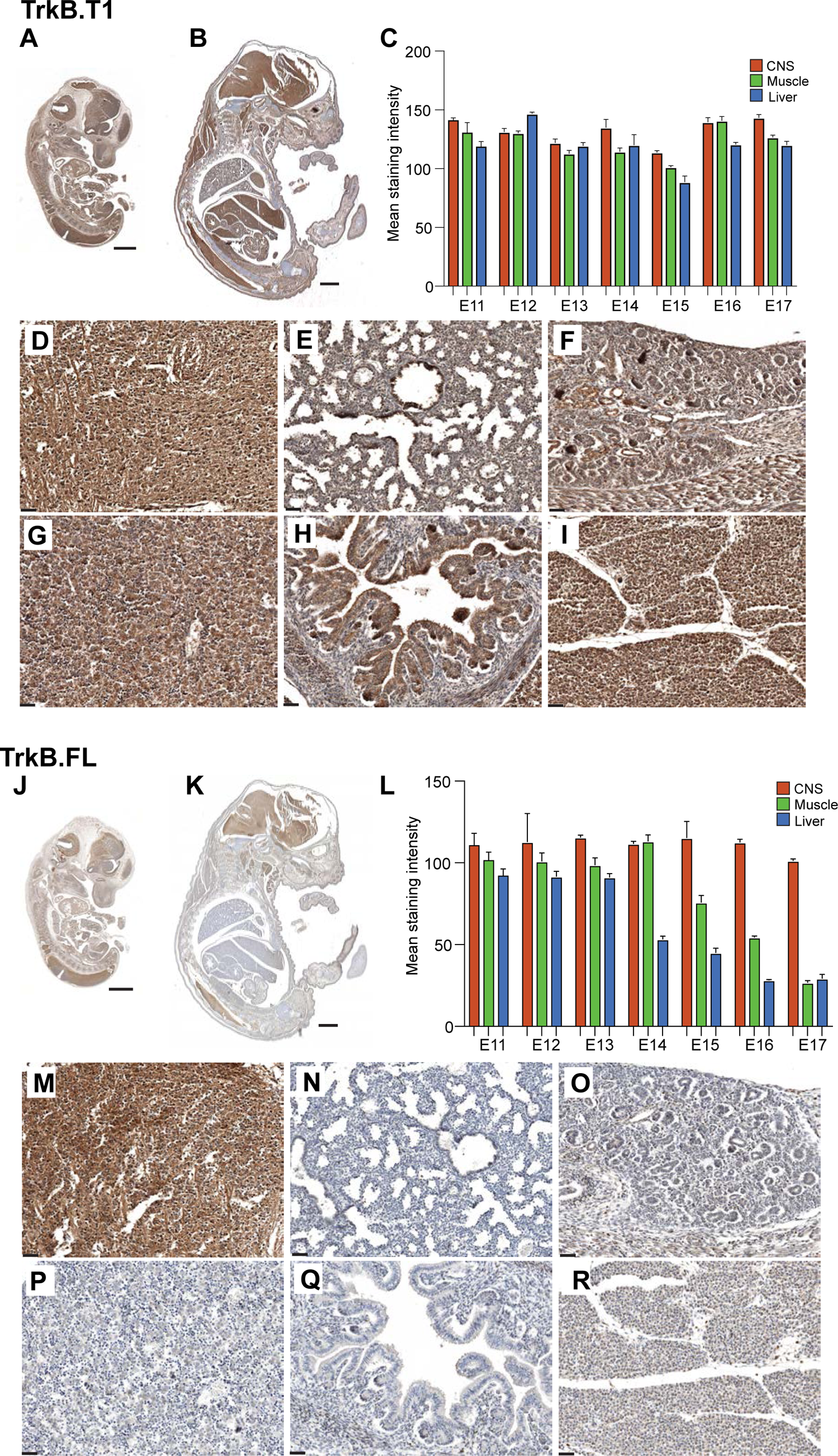
TrkB transcript expression across E10-E17 embryonic development. Representative micrographs of immunohistochemical staining of TrkB.T1 in the whole mount murine embryo section at E11**(A)** and E16 **(B)** Quantitative assessment of staining intensities in CNS (red), muscle (green) and liver (blue) at different developmental stages **(C)**. Representative micrographs for TrkB.T1 staining of **(D)** brain, **(E)** lung, **(F)** kidney, **(G)** liver, **(H)** small intestine, **(I)** skeletal muscle at E16. Immunohistochemical staining of TrkB.FL in the whole mount embryo section at E11**(J)** and E16 **(K)** with **(L)** quantitative assessment of staining intensities in CNS (red), muscle (green) and liver (blue) at different developmental stages. Representative micrographs for TrkB.FL staining in **(M)** brain, **(N)** lung, **(O)** kidney, **(P)** liver, **(Q)** small intestine, **(R)** skeletal muscle at E16. Scale bars: panels **A, B, J, K** = 1 mm; panels **D-L** and **M-R** = 100 μm.

### TrkB.T1 exerts significantly more transformation capacity compared to TrkB.FL in colony formation assay and is required for neurosphere formation

Given the widespread distribution of TrkB.T1 in multiple organ sites and cell types across embryonic development, and our recent findings that the TrkB.T1 splice variant predominates over the full-length kinase containing TrkB variant in gliomas and that this TrkB.T1 gene product enhances PDGF-driven glioma aggressiveness *in vivo* and amplifies PDGF-driven AKT and Stat signaling *in vitro* (Pattwell et al. 2020b), we wanted to compare the relative transformation potential of the kinase-containing TrkB.FL and kinase-deficient TrkB.T1 isoforms using a soft agar colony formation assay. Forced expression of the TrkB.T1 splice variant containing its unique 11-amino acid intracellular tail in 3T3 cells led to significantly enhanced colony formation (average colonies = 209) compared to overexpression of the kinase-containing variant, TrkB.FL (average colonies = 48), or control vector (average colonies = 19.33) (F (2,6) = 401.5, *p* <0.0001)(**Fig. 3A**). Using a short hairpin designed against TrkB.T1, we show that TrkB.T1 is necessary for neurosphere formation of primary *N/tv-a*;*Ink4a/Arf^-/-^* neural stem cells (NSCs) and that knocking down TrkB.T1 with RCAS-PDGFB-TrkB.T1 in primary NSCs leads to significantly less neurosphere formation compared to RCAS-PDGFB-shScrambled(SCR) control (**Fig. 3B; Supplemental Fig. S2B**);

**Figure 3:**
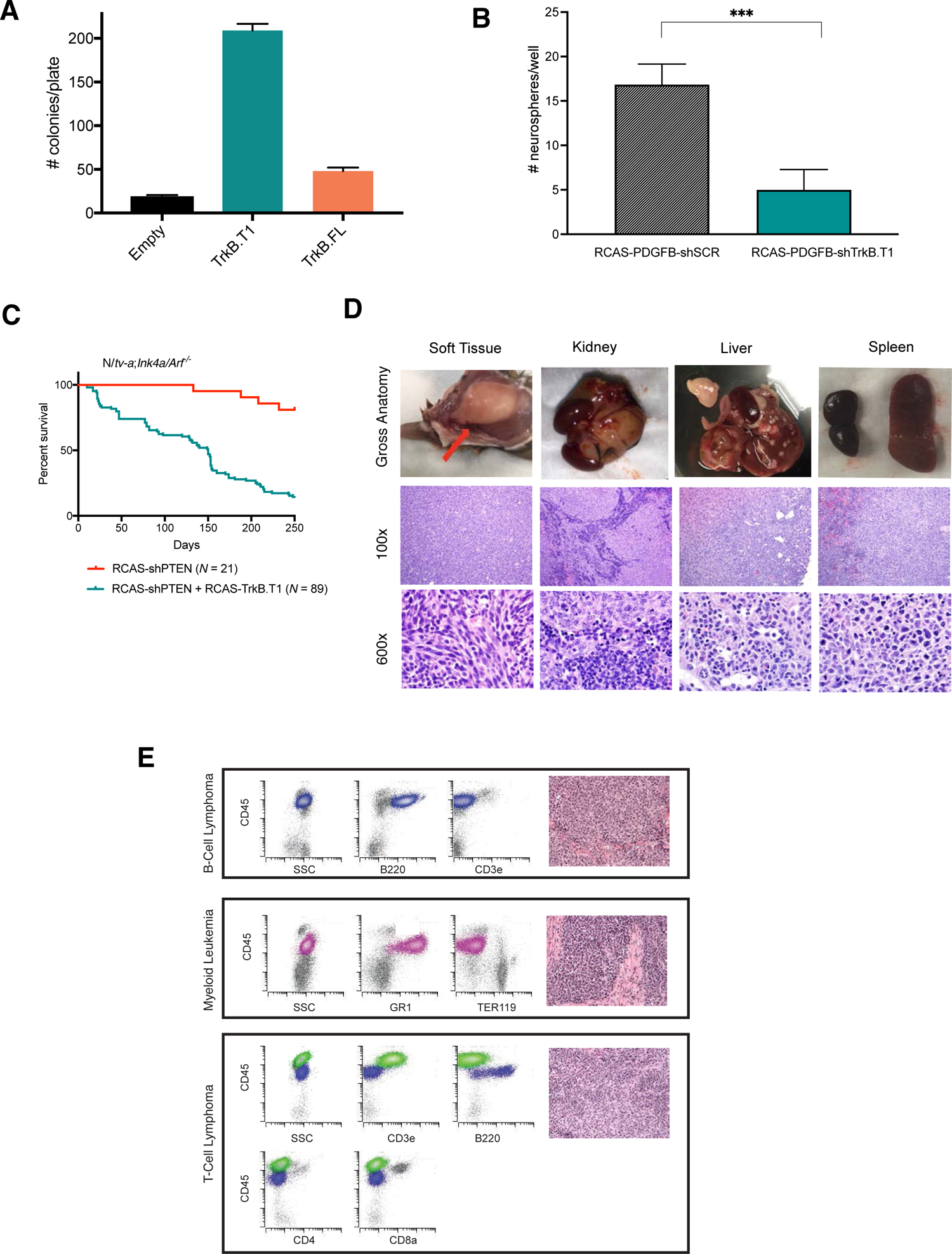
Gain and loss of function experiments show TrkB.T1 transforms 3T3 cells, is required for neurosphere formation, and causes solid and non-solid tumors throughout the body in mice when overexpressed in the context of tumor suppressor loss. (A) Soft agar colony formation assay in 3T3 cells shows significantly increased colonies with TrkB.T1 compared to control and TrkB.FL (*p* < 0.0001; graph represents mean + SEM). (B) TrkB.T1 knockdown with RCAS-PDGFB-shTrkB.T1 significantly decreases neurosphere formation in primary Nestin/*tv-a*;*Ink4a/Arf^-/-^* neural stem cells (RCAS-PDGFB-shTrkB.T1: M= 16.833 + SD 2.317 vs RCAS-PDGFB-shSCR: M = 5.000 + SD 2.280; t(5)=11.31, p <.0001). Survival curves for Nestin/*tv-a*;*Ink4a/Arf^-/-^* mice injected with RCAS-shPTEN alone or RCAS-shPTEN + RCAS-TrkB.T1 (C) demonstrate decreases in survival due to tumor burden outside the central nervous system for mice injected with RCAS-TrkB.T1 + RCAS-shPTEN vs. RCAS-shPTEN alone (median survival 131 days vs. 250 days, log rank hazard ratio 0.07243, 95% confidence interval 0.0.04768 to 0.11, *p* <0.0001) including soft tissue, kidney, liver, and spleen as shown in (D) and (E). Flow cytometry analysis of splenic tumor demonstrates a large B-cell lymphoma (CD45 bright/B220+; blue population). The histologic sections show a diffuse collection of large atypical cells with vesicular chromatin and high mitotic rate with a vague nodular pattern (20X). Flow cytometry analysis of splenic tumor demonstrates a myeloid leukemia (CD45+/GR1+; pink population) with extramedullary erythropoiesis (Ter119+/dim CD45; grey population). The histologic sections show focal collections of intermediate size cells with open chromatin and frequent mitosis, admixed with maturing erythroid cells (20X). Flow cytometry analysis of splenic tumor demonstrates an abnormal T-cell population (CD45 bright/CD3e+/CD4-/CD8a-; green population) consistent with a T-cell lymphoma admixed with reactive B-cells (CD45 bright/B220 variable; blue population). The histologic sections show large atypical cells with a high mitotic rate admixed with scattered small reactive lymphocytes and eosinophils (20X). ***p < 0.0001.

### Proteomic analysis reveals TrkB.T1-specific interactors with known developmental, oncogenic, and cell cycle signaling pathways

We explored potential interactors and upregulated pathways in an isoform specific NTRK2 manner in 3T3 cells separately transduced with pLJM1-lentiviral vectors containing either TrkB.T1-HA or TrkB.FL-HA (Pollard et al. 2006; Sun et al. 2008), in the presence or absence of serum. Lysates were collected and subjected to HA-tag pull down and subsequent affinity purification-mass spectrometry (AP-MS) proteomic analysis (Morris et al. 2014) to reveal potentially directly or indirectly interacting proteins (Morris et al. 2014).

Using HA-tagged *NTRK2* splice variants as bait, all immunoprecipitated proteins were identified, quantified, characterized, and subjected to Gene Ontology (GO) and Reactome Pathway enrichment analysis to understand and explore potential protein-protein interactions (**Fig. 4**). In the absence of serum, TrkB.FL expressing 3T3 cells began to die off, while TrkB.T1 cells continued to thrive, similar to the colony formation assay (**Fig. 3A**). For these serum starved conditions, 163 shared proteins were pulled down in complexes with both TrkB.T1 and TrkB.FL, 110 were pulled down specifically with TrkB.FL, and 720 proteins were pulled down specifically in complexes with TrkB.T1. Under normal culture conditions, in the presence of serum, 749 proteins were pulled down in complexes with both TrkB.T1 and TrkB.FL, 399 were pulled down specifically with TrkB.FL, and 565 proteins were pulled down specifically in complexes with TrkB.T1 (**Supplemental Dataset S1)**. GO and Reactome Pathway enrichment analyses on these clusters of affinity purified proteins revealed differential expression of a host of developmental, oncogenic, and mitotic pathway terms associated specifically with TrkB.T1 in the presence or absence of serum compared to the terms associated with TrkB.FL (**Supplemental** Fig. S3 **and Supplemental Dataset S1)**. Shared developmental and oncogenic terms between TrkB.T1 and TrkB.FL in the presence of serum implicated proteins or classes of proteins involved in FGFR2, IGFR, SLITs, ROBO, Hedgehog, TP53, and RUNX signaling, long known to be associated with neurotrophin signaling. Additional classes of proteins were found to be TrkB.T1 specific, under normal culture conditions and maintained in the absence of serum. Among these TrkB.T1-specific interactors (N=565) are proteins with known developmental and oncogenic roles, including Gli, Hedgehog, Wnt, TGF-ß, Ras, MAPK, MET, CaM, and other signaling pathways (**Fig. 4, Supplemental** Fig. S3, **and Supplemental Dataset S1)**. The distinct protein sets were pulled down with TrkB.T1 highlight a host of developmental, oncogenic, and cell cycle/cell division pathway binding partners that were not previously known to interact with TrkB.T1 along with several known interactors such as those associated with RhoA (Ohira et al. 2005; Ohira et al. 2006) and Ca^2+^ signaling (Rose et al. 2003).

**Figure 4:**
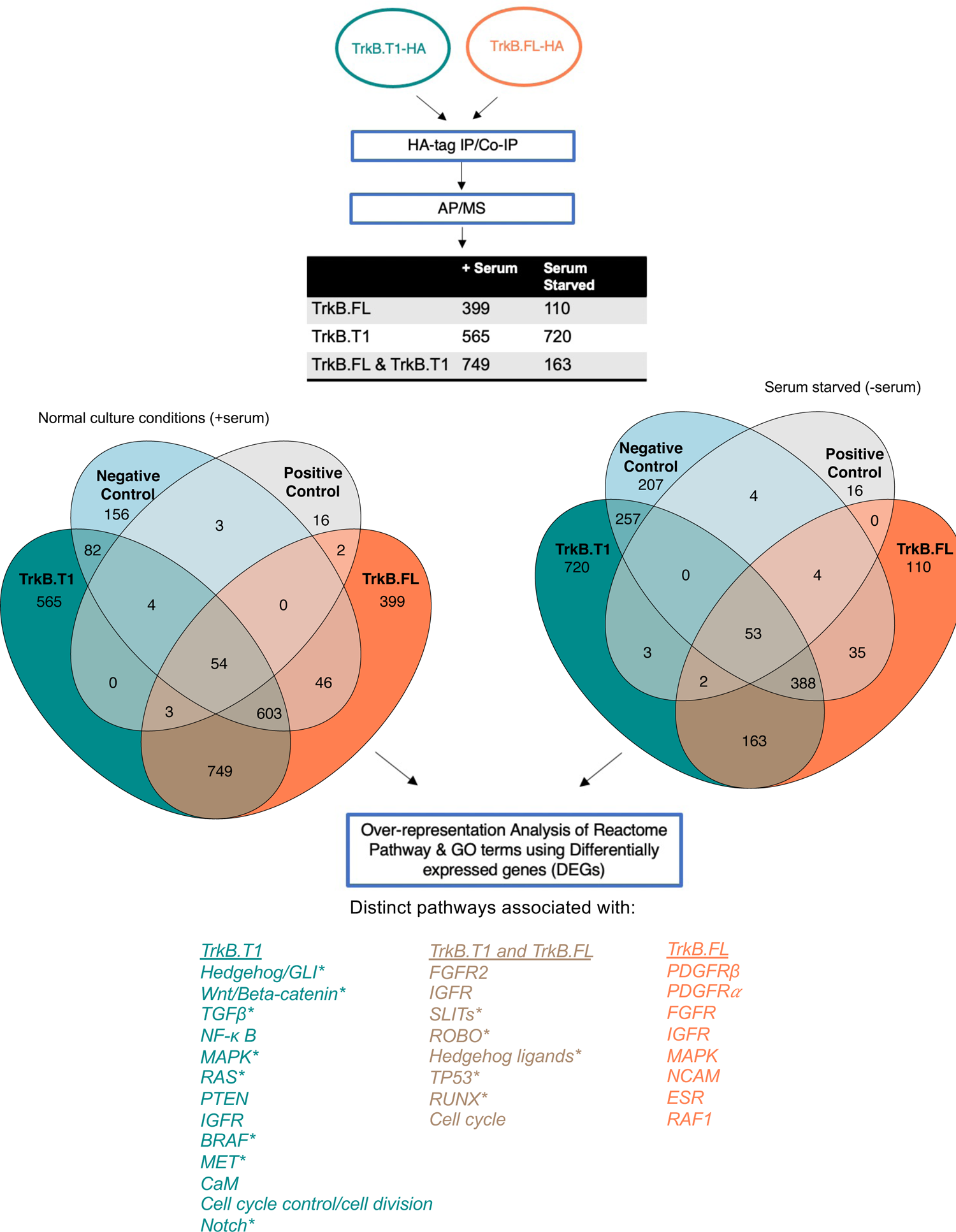
Affinity purification-mass (AP-MS) spectrometry reveals *NTRK2* splice variant interactors associated with developmental and oncogenic pathways. 3T3 cells were transduced with TrkB.T1-HA or TrkB.FL-HA, each in triplicate, and subjected to HA-pull down to identify direct and indirect interactors via AP-MS as shown in Supplemental Fig. S3 and Supplemental Dataset S1. Proteins specific to each isoform and shared between both isoforms (With serum: n=599 for TrkB.T1, n=399 for TrkB.FL, n=749 shared; Without serum: n=720 for TrkB.T1; n=110 for TrkB.FL, n=163 shared) were subjected to Gene Ontology (GO) and Reactome enrichment analysis to find proteins associated with developmental, oncogenic, and cell cycle/cell division pathways. *Developmental, oncogenic, and cell cycle/cell division associated signaling pathways listed are associated under normal culture conditions (with serum) unless shown with asterisk which denotes proteins associated with these pathways were found both in normal culture conditions and serum starved conditions.

### Forced postnatal expression of TrkB.T1 causes multiple cancer types in mice

Prior investigation into TrkB.T1’s role in gliomas revealed that a multitude of genes in PI3K/Akt/PIP/inositol phosphate pathways were significantly positively correlated with *NTRK2* expression in LGG and GBM compared to normal brain (Pattwell et al. 2020b) via Differential Gene Correlation Analysis (DCGA) suggesting a role for TrkB.T1 in the PI3K signaling pathway. Based on these results and the above developmental, transformation, and proteomic, data, we wanted to explore whether TrkB.T1 is capable of inducing tumors within and outside the CNS in the context of tumor suppressor loss. We chose mice with *CDKN2A* (Ink4a/Arf) loss and *PTEN* loss to explore in a sensitized genetic background as TrkB.T1 overexpression does not cause tumors on its own in wild-type mice (Pattwell et al. 2020a), while Pten is a negative regulator of Akt signaling and Ink4a/Arf loss is a frequent event in cancer. To answer this question, we used the RCAS/TV-A system, which allows for somatic expression of a gene of interest in particular cell types, to force overexpression TrkB.T1 and knockdown *PTEN* expression in nestin positive progenitor cells (Ozawa et al. 2014).

Mis-expression of factors or proteins that are benign or necessary in one cell type may lead to aberrant and non-canonical signaling in another cell type, as has been shown for Wnt signaling (Zhan et al. 2017). Because we saw TrkB.T1 interactions with other known developmental oncogenes in AP-MS and because nestin is expressed in a wide range of stem and progenitor cells during the development of various organ sites (not just neural and glial progenitors), this promoter was chosen to target RCAS delivery specifically to nestin positive cells. *N/tv-a*;*Ink4a/Arf^-/-^* mice were injected intraperitoneally (i.p.) or intramuscularly (i.m.) with RCAS-TrkB.T1 and RCAS-shPTEN. While neither RCAS-shPTEN nor RCAS-TrkB.T1 alone caused cancer in this genetic background during the time course of this experiment, the combination of RCAS-shPTEN and RCAS-TrkB.T1 injections caused a range of solid and non-solid tumors originating from nestin-positive cells outside of CNS (**Fig. 3C**). These included soft tissue sarcomas, carcinomas arising in the kidney and lymphoid leukemias and lymphomas (**Fig. 3D, 3E**). Immunohistochemistry on these solid tumors confirmed that these tumors throughout the body had both high levels of TrkB.T1 and loss of PTEN expression (**Supplemental Fig. S4A – S4D**) demonstrating that forced expression of TrkB.T1 in combination with PTEN loss has the potential to form tumors in the same organ sites that once expressed TrkB.T1 embryonically.

### TrkB.T1 is the predominantly expressed form of TrkB RNA and protein in adult and pediatric tumors

The large number and breadth of murine tumor types induced by TrkB.T1 expression, combined with the knowledge that in gliomas the predominant form of TrkB is the TrkB.T1 splice variant, prompted an in-depth pan-cancer analysis of *NTRK2* transcript expression across all organ sites within The Cancer Genome Atlas (TCGA). Similar to what has been reported for gliomas (Pattwell et al. 2020b), expression of the full-length, kinase containing TrkB.FL is minimal to absent across the majority of tumor types within TCGA, while TrkB.T1 expression is consistently high (**Fig. 5, Supplemental Dataset S2**). In depth transcript characterization of all TCGA tumors harboring known NTRK fusions recapitulates this pattern and shows that regardless of TRK fusion (NTRK1, NTRK2, NTRK3), expression of the TrkB.T1-specifiic exon (encoding for the unique 11 amino acid intracellular region) remains consistently higher than all kinase containing TRK exons, regardless of the particular TRK fusion or tumor type (**Supplemental** Fig. S5, **Supplemental Dataset S3**).

**Figure 5:**
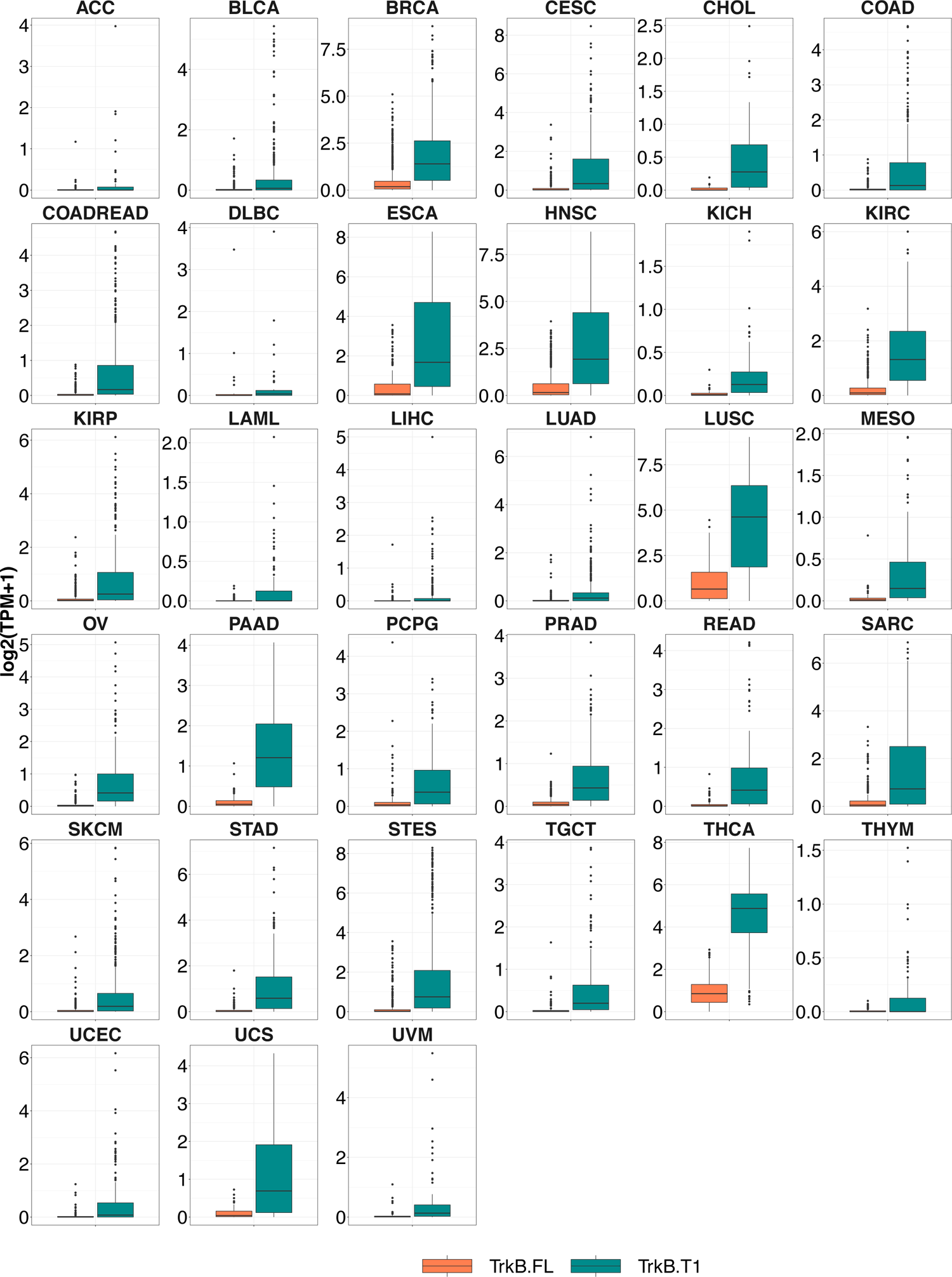
Pan-cancer *NTRK2* Transcript Analyses. *NTRK2* transcript analysis shows minimal to zero expression of TrkB.FL transcript and increased expression for TrkB.T1 transcript across all TCGA organ sites: adrenocortical carcinoma (ACC), bladder urothelial cancer (BLCA), breast invasive carcinoma (BRCA), cervical squamous cell carcinoma and endocervical adenocarcinoma (CESC), cholangiocarcinoma (CHOL), colon adenocarcinoma (COAD), colorectal adenocarcinoma (COAD/READ), lymphoid neoplasm diffuse B-cell lymphoma (DLBC), esophageal carcinoma (ESCA), head & neck squamous carcinoma (HNSC), kidney chromophobe (KICH), kidney renal clear cell carcinoma (KIRC), kidney renal papillary cell carcinoma (KIRP), acute myeloid leukemia (LAML), liver hepatocellular carcinoma (LIHC), lung adenocarcinoma (LUAD), lung squamous cell carcinoma (LUSC), mesothelioma (MES), ovarian serous cystadenocarcinoma (OV), pancreatic adenocarcinoma (PAAD), pheochromocytoma and paraganglioma (PCPG), prostate adenocarcinoma (PRAD), rectum adenocarcinoma (READ), sarcoma (SARC), skin cutaneous melanoma (SKCM), stomach adenocarcinoma (STAD), stomach and esophageal (STES), testicular germ cell tumor (TGCT), thyroid carcinoma (THCA), thymoma (THYM), uterine corpus endometrial carcinoma (UCEC), uterine carcinoma (UCS), uveal melanoma (UVM). Data are represented as boxplots where the middle line is the median, the lower and upper hinges correspond to the first and third quartiles (the 25th and 75th percentiles), the upper whisker extends from the hinge to the largest value no further than 1.5 * IQR from the hinge (where IQR is the inter-quartile range, or distance between the first and third quartiles) and the lower whisker extends from the hinge to the smallest value at most 1.5 * IQR of the hinge while data beyond the end of the whiskers are outlying points that are plotted individually.

In addition to the data from TCGA, which is comprised of mostly adult tumors, previous studies have also shown a potential role for TrkB in pediatric tumors and its gene products have been shown to be expressed at high levels in high-risk neuroblastoma (NBL) and correlated with poor prognosis (Geiger and Peeper 2005; Nakamura et al. 2014) while levels of TrkB gene products have been shown to be correlated with unfavorable outcome in Wilms tumor (WT) patients (Eggert et al. 2001). Transcript specific analysis of pediatric data from the Therapeutically Applicable Research to Generate Effective Treatments (TARGET) program confirms that, similar to adult tumors in TCGA, TrkB.T1 expression is the predominant isoform compared to TrkB.FL in several cancer types, including Wilms tumor (WT), rhabdoid tumor (RT), neuroblastoma (NBL), and clear cell sarcoma of the kidney (CCSK) (**Supplemental** Fig. S6**, Supplemental Dataset S2**).

To extend upon these transcript results demonstrating predominant TrkB.T1 expression across a wide range of human tumors, we next determined how well RNA transcripts correlate with TrkB.T1 protein distribution. Immunohistochemistry on a tissue microarray containing a wide assortment of human tumor types (**Fig. 6**) reveals variable, increased TrkB.T1 expression and distribution, with high TrkB.T1 H-score values (**Supplemental Dataset S4**) for nearly all tumor types. These results show that both RNA transcript expression and protein levels for TrkB.T1 are high across all tumor types in available datasets, suggesting that this is the *NTRK2* splicing choice made across cancers and predominates in non-neuronal cells of the embryo as well.

**Figure 6.**
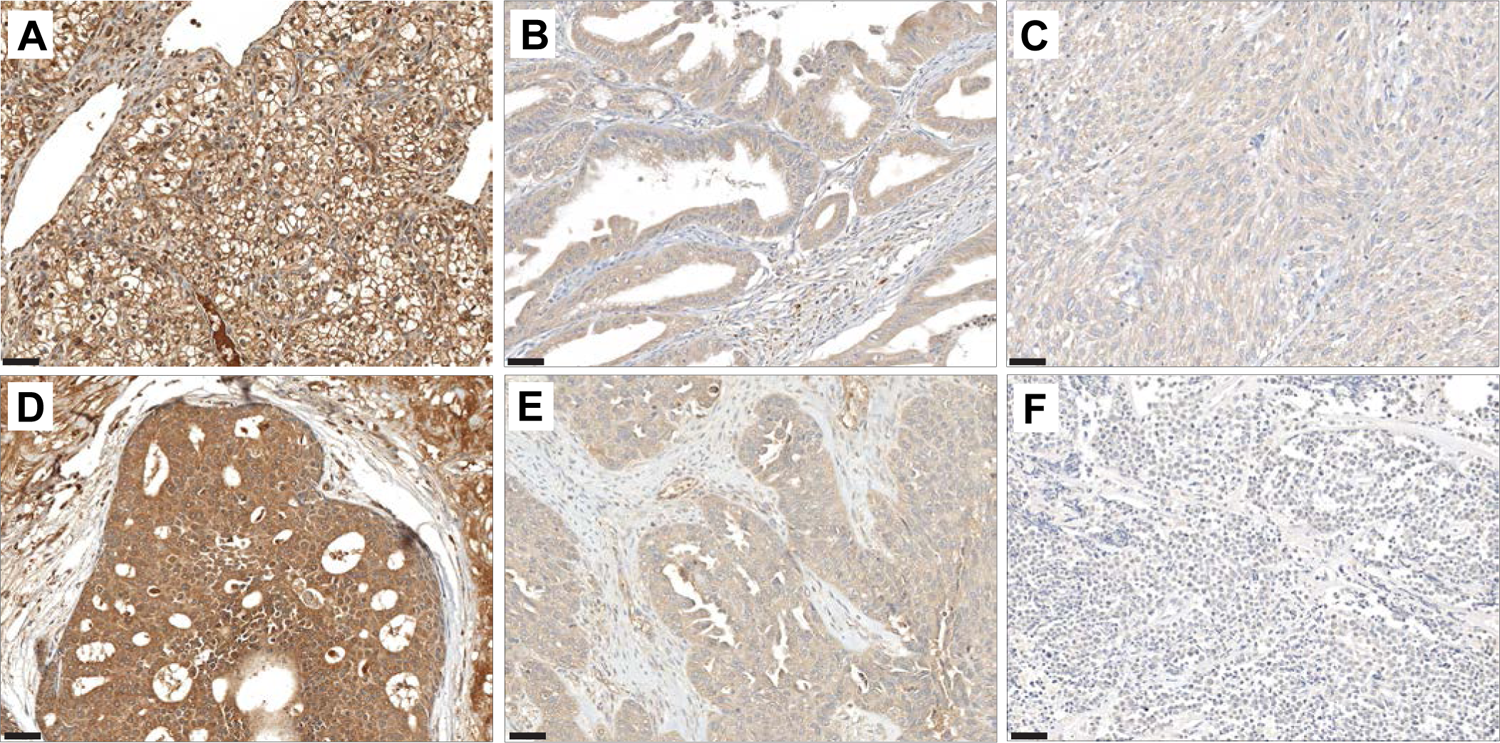
TrkB.T1 has variable expression across multiple human tumor types. Representative micrographs of TrkB.T1 expression in **(A)** clear cell renal cell carcinoma**, (B)** colonic adenocarcinoma, **(C)** leiomyosarcoma**, (D)** mammary carcinoma (ductal carcinoma in situ is shown here), **(E)** pancreatic adenocarcinoma, and **(F)** seminoma. Scalebar indicates 100 μm.

Taken together, these data show that TrkB.T1 is the *NTRK2* isoform expressed in multiple cell types across embryonic development, is expressed at surprisingly high rates across adult and pediatric cancers from multiple sites within TCGA and TARGET (**Fig. 5, Supplemental** Fig. S6**, Supplemental Dataset S2-S4)** and is capable of transforming cells in vitro and causing tumors across various organ sites in the context of loss of the common tumor suppressors Ink4a/Arf and PTEN in mice (**Fig. 3, Supplemental Fig. S4**), potentially through its interactions with known oncogenic and developmental signaling pathways (**Supplemental Fig. S3, Supplemental Dataset S1**). These data highlight the TrkB.T1 splice variant as one candidate whose aberrant expression with concurrent tumor suppressor loss causes cancer in the same developed organs where it played a role in early embryonic development.

## Discussion

While many point mutations have been discovered in cancer, only a small majority of these mutations have been shown to be causal. These rare driver mutations contribute to oncogenic signaling by altering pathways that are otherwise tightly regulated in normal tissues (Simanshu et al. 2017). It is known that cancer cells exhibit vast transcriptomic alterations and that cancer specific isoforms are not merely byproducts of abnormal physiology (El Marabti and Younis 2018), yet most cancer associated splicing choices, like point mutations, are not oncogenic drivers.

Differential splicing choices allow for highly regulated expression of specific transcripts in normal development, yet expression of specific splice variants at developmentally inappropriate timepoints or in the wrong cell types may be detrimental. While it is known that aberrant splicing mechanisms exist in cancer (Oltean and Bates 2014; Sveen et al. 2016; Escobar-Hoyos et al. 2019), in this study, we show that a splice variant with a physiologic role in early organogenesis, but whose expression is not found in corresponding mature tissue, can be a driver of cancer. TrkB.T1 appears to be a candidate whose regulated splicing choice appears in normal embryonic development and whose mis-expression at later postnatal time points is oncogenic.

While the embryogenic data offers correlative insight, the causal RCAS/tv-a mouse studies show that aberrant expression of TrkB.T1 causes neoplasms at multiple primary sites, potentially by trapping cells in an undifferentiated state similar to their early embryogenic stage via interactions with known signaling pathways such as Wnt, Hedgehog, TGF-ß, or Ras. In the embryonic sci-RNA-Seq3 data, TrkB.FL and TrkB.T1 are rarely, if ever, expressed in the same cell and they also have a unique set of interactors as evidenced by HA affinity purification mass spectrometry, suggesting that TrkB.T1 does not function through transactivation of TrkB.FL, but potentially through other signaling pathways. These data highlight the TrkB.T1 splice variant as a promising candidate for future studies investigating the role of *NTRK2* in development and oncogenesis across a range of organ sites and tumor models beyond the known neurobiological roles of TrkB.T1(Dorsey et al. 2006; Carim-Todd et al. 2009; Fulgenzi et al. 2015) and further strengthen the potential links between the tightly regulated splicing patterns of development with the aberrant splicing patterns observed in cancer. Approaches to studying splicing factors and alternatively spliced transcripts such as those delineated here should not be restricted to particular genes, stages of development, or organ sites and may uncover promising new avenues for diagnostics or therapeutics by revealing additional developmentally regulated, oncogenic splice variants.

## Acknowledgements

We thank James Yan, Jenny Zhang, Deby Kumasaka, and Denis Adair for continued technical and administrative assistance and support throughout these experiments. We thank Francis S. Lee at Weill Cornell Medical College for advising on experiments and data. We thank Phil Gafken and Lisa Jones for help in carrying out IP-MS experiments.

## Author Contributions

S.S.P. proposed the scientific concepts, designed and performed experiments, generated data, assisted with bioinformatic analyses, and wrote the manuscript. S.A., H.B., N.N. performed bioinformatic analyses, designed bioinformatic pipelines, and generated data. S.A., N.N., M.Z. helped with data visualization. J.C. and J.S. contributed to the single cell data and provided sci-RNA-seq3 expertise. P.J.C. and M.C.H. examined tissue slides for human and mouse, generated photomicrographs, offered neuropathology and general pathology expertise and contributed to the manuscript. T.O. provided necessary reagents, supervised initial experimental design, and offered technical guidance throughout. K.R.L. performed experiments, generated data, and provided hematopoietic insight. N.H. and D.A.B. contributed to transformation and NSC experiments and collected data. M.J.W. helped with interpretation of sarcoma related data and manuscript preparation. F.S., V.V.P., and D.A.B. assisted with design, generation, and validation of RCAS vectors. N.R-L. helped with mass spectrometry experiments and subsequent data analysis. E.C.H. contributed to the overall experimental design, supervised the project, and offered critical feedback throughout the project and manuscript revisions. All authors discussed data and contributed to the final manuscript.

## Data and materials availability

The data and materials that support the findings of this study are included with the manuscript and supplemental data files and also available from the corresponding author upon reasonable request. All R scripts used in this manuscript are freely available via: https://github.com/sonali-bioc/Pattwell_transcript_sci-RNA-seq3

**Supplemental Dataset S1-S4**

**Supplemental Figures S1-S6** https://atlas.fredhutch.org/fredhutch/ntrk2/

## Methods

### Bioinformatic Analysis

#### Quantification of Transcript data from sci-RNA-seq3

SAM alignment files for the MOCA dataset were downloaded from https://shendure-web.gs.washington.edu/content/members/cao1025/public/nobackup/. The “cell_annotation.csv” which contained *t*-SNE coordinates, UMAP coordinates, and information about clusters and trajectories was downloaded from https://oncoscape.v3.sttrcancer.org/atlas.gs.washington.edu.mouse.rna/downloads. The “Comprehensive gene annotation” file was downloaded from https://www.gencodegenes.org/mouse/release_M12.html For each of the 2,062,641 cells, we calculated the number of strand-specific UMIs for each cell mapping to transcripts of each gene with the Python v.2.7.13 HTseq (Anders et al. 2015) package using GENCODE vM12 (Harrow et al. 2012). Monocle3 (Trapnell et al. 2014) was used to construct a cell_data_set object with all the transcript expression data, preprocess_cds() was used to normalize the transcript expression. The expression of TrkB.T1 and TrkB.FL were visualized over existing UMAPs and *t*-SNEs provided by Cao et al using ggplot2 (Wickham 2009). An interactive website (https://atlas.fredhutch.org/fredhutch/ntrk2/) has been put together to facilitate further exploration of the NTRK2 transcript data across various trajectories and cell types.

#### Obtaining & Transforming Transcript Level Data from TCGA and TARGET

Transcript Data (TCGA RNAseqV2 RSEM data) for TCGA and TARGET organ sites was downloaded from Broad’s FireBrowse website (https://gdac.broadinstitute.org/). Transcript data (TPM data) was also downloaded for TARGET from UCSC Xena: (https://xenabrowser.net/datapages/?dataset=target_RSEM_gene_tpm&host= https%3A%2F%2Ftoil.xenahubs.net&removeHub= https%3A%2F%2Fxena.treehouse.gi.ucsc.edu%3A443). The results published here are in whole or part based upon data generated by TCGA (Cancer Genome Atlas Research et al. 2013) and the Therapeutically Applicable Research to Generate Effective Treatments (https://ocg.cancer.gov/programs/target) initiative, phs000218. The data used for this analysis are available at https://portal.gdc.cancer.gov/projects.”

To make the RSEM data from TCGA and TARGET, and TPM data comparable, we converted RSEM counts from TCGA to TPM counts using the following formula (Li and Dewey 2011): RSEM can be multiplied by 10^6^. For transcript analyses, NTRK2 transcript IDs were manually aligned to confirm sequence homology and are as follows: TrkB.FL (UCSC: uc004aoa.1; Ensembl: ENST00000376213.1,NTRK2_201), TrkB.T1 (UCSC: uc004aob.1; Ensembl: ENST00000395882.1, NTRK2_204). Transcript Data was visualized using boxplots using R package ggplot2 (Wickham 2009). Gene fusions for all TCGA fusions was obtained from GDC portal (https://portal.gdc.cancer.gov/). The transcript levels for TrkB.T1 and N-terminal truncated TrkB.T1, both containing the 11 amino acid intracellular region, for TCGA samples ids containing TRK gene fusions (including fusions in NTRK1, NTRK2 and NTRK3) were compared to the mean of all other NTRK transcripts.

#### Affinity-purification mass spectrometry analysis

GO and Reactome Pathway enrichment analysis for genes associated with enriched mass spectrometry proteins (Fabregat et al. 2018; Jassal et al. 2020) was done using R Bioconductor Packages clusterProfiler v 3.4.4 (Yu et al. 2012) and dot plots were made using R Bioconductor package DOSE.

### Generation of murine tumors

The RCAS/tv-a system used in this work has been described previously for murine tumor modeling in immunocompetent mice(Holland et al. 2000). DF-1 cells were transfected with the relevant RCAS viral plasmids using Extreme-Gene 9Transfection reagent (Roche) accordingly to manufacturer’s protocol. The cells were maintained for three passages, as described above to ensure viral propagation to all cells. After confirmation of RCAS-inserts by western blot, DF1s (passage 4 or later) were used for injection into murine brain. Newborn *N/tv-a*;*Ink4a/Arf^-/-^* pups (P0-P1) were injected (Hamilton syringe #84877) with 1 μL of approximately 1×10^5^ DF-1 cells infected with and producing relevant RCAS viruses suspended in serum-free DMEM (TrkB.T1 and shPTEN) (Ozawa et al. 2014). Simultaneous delivery of two RCAS viruses was performed by the injection of 1μL of approximately 2 x 10^5^ DF-1 cells mixed with equal ratio. Mice were monitored for the duration of the study (250 days) to check for tumor related symptoms such as palpable masses, lethargy, weight loss, seizure, hyperactivity, altered gait, poor grooming, macrocephaly, paralysis. Mice with severe hydrocephalus presumably due to injection trauma or an inflammatory response against the DF-1 cells were excluded from survival analysis in this study. All animal experiments were approved by and conducted in accordance with the Institutional Animal Care and Use Committee of Fred Hutchinson Cancer Research Center (protocol #50842).

### Mouse Tissue Processing

Mouse tissue (including normal brains, tumor bearing brains, solid tumors) were removed, fixed in 10% neutral-buffered formalin for a minimum of 72 hours and embedded into paraffin blocks. 5µm serial sections were cut from formalin-fixed paraffin embedded specimens and mounted on slides.

### Immunohistochemistry

Immunohistochemical staining was performed on 5um formalin-fixed/paraffin-embedded tissue sections using a Discovery XT Ventana Automated Stainer (Ventana Medical Systems, Inc)., run using standard Ventana reagents a and Vector secondaries for staining with the TrkB.T1 SPEH1_D12 scFv-Fc fusion (Pattwell et al. 2020b) at 1:500 and PTEN (Cell Signaling #9188, Lot #4) at 1:100 or TrkB (kinase specific against amino acid 810; abcam #ab18987, lot #GR3280550-2) at 1:250. All embryonic histology slides and human tissue microarray (BioMax, U.S. #BC001134b) slides were scanned using a Ventana DP200 slide scanning system (Roche Diagnostics) at 20x magnification. Digital images of different organ during mouse development were analyzed using the Fiji image analysis software as described previously (Schindelin et al. 2012; Haffner et al. 2017). Staining intensities in human tumor tissues were assessed using a semi-quantitative H-score system by multiplying the intensity of the stain (0: no staining; 1: weak staining; 2: moderate staining; 3: intense staining) by the percentage (0 to 100) of cells showing that staining intensity (H-score range, 0 to 300) as described previously(Haffner et al. 2011).

### Flow cytometry

Rodent tumors were harvested and resuspended as single cells in PBS/0.3% BSA. Cells were washed and incubated with the following antibodies: CD45-APC-Cy7 (Biolegend, clone 30-F11, catalog #103116, lot #B242535), CD3e-PE (BD, clone 145-2C11, catalog #553061, lot #22126), GR1-PerCP (BD, clone Ly-6G/Ly-6C, catalog #552093, lot #73108), TER119-APC (BD, clone Ter-119, catalog #557909, lot #42622), B220-Alexa 647 (BD, clone RA3-6B2, catalog #557683, lot #22218), CD4-PerCP (BD, clone L3T3, catalog #553654, lot #60912), and CD8a-FITC (BD, clone Ly-2, catalog #553030, lot #46675), as indicated. Cells were analyzed on a custom built LSR II flow cytometer (BD). Data compensation and analysis were performed by using noncommercial software developed in our laboratory (Wood 2006).

### Soft agar colony formation assay

#### Lentiviral Production

293T packaging cells were seeded in 100mm plates at a density of 3.8 x 10^6^ cells per plate. 293T cells were supplemented with complete DMEM media (Dulbecco’s Modified Eagle Media (ThermoScientific, catalog #11966025) containing 10% Fetal Bovine Serum (Fetal Bovine Serum (ThermoScientific, catalog #26140079) and 5% Penicillin/Streptomycin. 293T cells were split 3 times a week in order to maintain healthy density, and incubated at 37°C. Once the optimal number of 293T 10mm plates were produced, they were transfected with psPAX2 (Addgene, catalog #12260), pMD2.G (Addgene, catalog #12259), and the gene of interest containing pLJM1 lentiviral packaging plasmids (Addgene, catalog #91980) at concentrations of 1.3 pmol, 0.72 pmol, and 1.64 pmol. 293T cells were transfected using the Thermofisher Lipofectamine 3000 transfection protocol (ThermoFisher, catalog # L3000001). The media was replaced after 18 hours in order to permit lentiviral production in transfection reagent free media.

Viral media was harvested after 48, 72, and 96 hours and replaced with fresh media after each harvest. The viral supernatant was filtered through 0.45um syringe filters Millex-HP Syringe Filter Unit 0.45 µm (Millipore Sigma, SLHP033RS) to remove cellular debris and other contaminants.

NIH-3T3 cells were seeded in 100mm plates at a density of 2.2 x10^6^ cells per plate, and supplemented with complete DMEM containing 10% calf serum (Calf Serum (ThermoScientific. catalog#16170086) and 5% Penicillin/Streptomycin. Once the NIH-3T3 plates reached optimal confluency, NIH-3T3 cells were supplemented with the viral harvest media and left to incubate for 24 hours. The viral harvest was then removed and fresh complete DMEM media with 2µg/ml of puromycin. The NIH-3T3 cells were incubated for a week under these selection conditions until the population density rebounded to plate confluency.

#### Soft Agar Colony Formation Assay

Once the NIH-3T3 cells under each condition were confluent, we initiated the Soft Agar Colony Formation assay. A 3% agarose solution (Agarose; Guidechem, catalog number: 9012-36-6) was microwaved for 1 minute and stored in a 45°C water bath in order to maintain its liquid state. The 3% agarose solution was aliquoted and diluted with complete DMEM media to yield a 0.6% agarose/media solution. This new mixture was poured into 6-well plates at 1ml and allowed to solidify for 30 min. Aliquots of 0.6% solution were then diluted with the NIH-3T3 cell media suspension so that the solution was composed of 0.3% agarose and contained a concentration of 2 x 10^3^ cells/ml. This solution was stored at 37°C rather than 45°C to prevent cell death. The solution was plated atop the now solid 0.6% agarose layer and allowed to solidify at room temperature for 1 hour.

The agarose suspension of cells was incubated at 37°C for 30 days while supplemented with a 1ml feeder layer of DMEM with 1ug/ml of puromycin and 10% calf serum. After 30 days, the feeder layer was removed and a solution of 0.005% crystal violet dye was plated onto the agarose to darken the formed colonies. Colonies were photographed at 10X and 20X magnification, and were counted at 4X magnification.

### shTrkB.T1 and neurosphere formation

#### Real-time PCR

RNA from Nestin(N)/tv-a;Ink4a/Arf^-/-^ neural progenitors was isolated using the RNeasy Kit (Qiagen, Catalog# 74104) and 1μg of total RNA was reverse transcribed using the High-Capacity cDNA Reverse Transcription Kit (Applied Biosystems, Catalog # 4368814). For quantitative analysis, we utilized the PowerUP SYBR Green Master Mix (Applied Biosystems, Catalog #A25742) following the manufacturer’s instructions. Quantitative PCR was performed using the QuantStudio 7 Real-Time PCR system. We have utilized previously published primers (Fulgenzi et al. 2015) for TrkB.T1 (Forward: AGCAATCGGGAGCATCTCT and Reverse: TACCCATCCAGTGGGATCTT) and TrkB.FL (Forward: AGCAATCGGGAGCATCTCT and Reverse: CTGGCAGAGTCATCGTCGT). The threshold cycle number for the genes analyzed was normalized to Actin (Forward: CGTGGGCCGCCCTAGGCACCA and Reverse: CTTAGGGTTCAGGGGGGC) and TrkB.FL and TrkB.T1 levels were normalized to uninfected (N)/tv-a;Ink4a/Arf^-/-^ neural progenitors.

#### Neurosphere formation

Neural progenitors isolated from (N)/tv-a;Ink4a/Arf^-/-^ pups at P0 as previously described (Pattwell et al. 2020a) were infected with RCAS-PDGFB-shSCR (scrambled control) or RCAS-PDGFB-shRNA against TrkB.T1, utilizing an RCAS/tv-a system as has been described previously described (Holland et al. 2000; Ozawa et al. 2014).

The specific hairpins against TrkB.T1 used in are:

shTrkB.T1#1: (GAAAAAGCTAACCACCTGCCCTTTAGATCTCTTGAATCTAAAGGGCAGGTGGTTA GCGGG),

shTrkB.T1#2: (GAAAAAGCACTCTCCTCCGCTTTATCTTCTCTTGAAAGATAAAGCGGAGGAGAGTGCGGGGATC),

shTrkB.T1#3: (GAAAAAGTCATAAGATCCCCCTGGATTCTCTTGAAATCCAGGGGGATCTTATGAC GGG), along with the scrambled control: (GAAAAAGCTCTACAACCGCTCATCATATCTCTTGAATATGATGAGCGGTTGTAGAG CGGG).

Briefly, RCAS virus was produced in Chicken fibroblast (DF1) cells maintained with 10% fetal bovine serum (Clonetech, Catalog# 631101) in Dulbecco’s modified Eagle medium (Thermo Scientific, Catalog# 11995-073) with 1% Penicillin/Streptomycin (Fisher Scientific, Catalog# 30-002-CI) at 39 degrees Celsius. DF1 cells were transfected with the indicated RCAS plasmid using X-tremeGENE 9 DNA transfection reagent (Roche, Catalog# 06365809001) according to the manufacturer’s protocol. RCAS transgene expression was confirmed via qPCR analysis. DF1 cells were then incubated with murine neurosphere media for 24 hours to package the indicated RCAS plasmid into virus. Our murine neurosphere media consisted of 45 ml Neurocult Mouse Basal media (Stem Cell Technologies, Catalog# 05700), 5 ml of mouse Neurocult Proliferation Supplement (Stem Cell Technologies, Catalog# 05701), 50 μL of 20 μg/ml EGF (Peprotech Inc, Catalog# 100-47), 50 μL of 10 μg/mL bFGF (Peprotech Inc, Catalog# 100-18B), 125 μL Heparin (Stem Cell Technologies, Catalog# 07980), 500 μL P/S (Fisher Scientific, Catalog# 30-002-CI). This conditioned media was then diluted 1:1 in fresh murine neurosphere media and used to incubate (N)/tv-a Ink4a^-/-^ Arf^-/-^ neural progenitor cells overnight. (N)/tv-a Ink4a^-/-^ Arf^-/-^ neural progenitor cells were then washed with PBS and maintained in fresh neurosphere media for 48 hours. They were then seeded at a concentration of 50,000 cells/well in ultralow attachment 6-well plates (Corning, Catalog# 3471). Fresh neurosphere media (0.5mL/well) was added once a week. Neurospheres of >100 μm in diameter were manually counted using a Leica DMIL LED microscope with an eyepiece graticule at 7, 14 and 21 days, data displayed for 21 days.

### Affinity Purification Mass Spectrometry (AP/MS)

Similar to methods for the colony formation assay, pLJM1 (Addgene) constructs containing the NTRK2 inserts of interest (TrkB.FL, TrkB.T1, or GFP) were transfected into 293T cells, along with psPAX and pMD2.G packaging plasmids (Addgene), using polyethylenimine (Polysciences). Fresh media was added 24 hours later and viral supernatant harvested 24 hours after that. For infection of 3T3 cells, 1e5 cells/well were seeded into 6-well plates. Lentivirus was used unconcentrated and cells were infected at a MOI<1 24 hours after seeding. 72 hours after seeding, selection was begun for cells successfully expressing the constructs using 2 μg/mL puromycin (for 7 days).

3T3 cells were expanded after selection to create a stable line and collected for HA-tag Co-IP analysis (Thermo Scientific™ Pierce™ HA-Tag IP/Co-IP Kit; Cat #26180) in M-Per buffer (M-PER™ Mammalian Protein Extraction Reagent; Cat #78501) three weeks post infection. This kit includes immobilized antibody resin and improved protocols over other methods with a GST-PI3K-SH2-HA fusion protein as positive control, is robust, specific, and has elution conditions compatible with subsequent downstream analysis. To remove detergents needed for Co-IP prior to downstream MS analysis, proteins were resolved by SDS/PAGE (NuPAGE 10% Bis/Tris; LifeTech) according to XCell Sure Lock™ Mini-Cell guidelines. Gels were stained with Simply Safe Blue and bands were excised and submitted to the Fred Hutch Proteomics Core for MS.

Gel slices were washed with water, 50% acetonitrile/50% water, acetonitrile, ammonium bicarbonate (100 mM), followed by 50% acetonitrile/50% ammonium bicarbonate (100 mM). The solution was removed, and the gel slices were dried in a speed vac. The gel slices were reduced with dithiothreitol (10 mM in 100 mM ammonium bicarbonate) at 56 °C for 45 min. The solution was removed and discarded. The gel slices were alkylated with 2-chloroacetamide (55 mM in 100 mM ammonium bicarbonate) and incubated in the dark at ambient temperature for 30 min. The solution was removed and discarded. The gel slices were washed with ammonium bicarbonate (100 mM) for 10 min on a shaker and an equal amount of acetonitrile was added and continued to wash for 10 min on a shaker. The solution was removed, discarded and the gel slices were dried in a speed vac for 45 min. The gel slices were cooled on ice and a cold solution of trypsin (Promega, Madison, WI) 12.5 ng/μL, in ammonium bicarbonate (100 mM) was added, enough to cover the gel slice. After 45 min, the trypsin solution was removed, discarded and an equal amount of ammonium bicarbonate (50 mM) was added and incubated overnight at 37 °C with mixing. Samples were spun down in a microfuge and the supernatants were collected. Peptides were extracted from the gel slices by adding 0.1% trifluoroacetic acid (TFA) enough to cover the slices and mixed at ambient temperature for 30 min. An equal amount of acetonitrile was added, and the samples were mixed for an additional 30 min. The samples were spun on a microfuge and the supernatants were pooled. The supernatants were concentrated in a speed vac. All samples were desalted using ZipTip C_18_ (Millipore, Billerica, MA) and eluted with 70% acetonitrile/0.1% TFA. The desalted material was concentrated in a speed vac.

The generated peptide samples were brought up in 2% acetonitrile in 0.1% formic acid (20 μL) and analyzed (18 μL) by LC/ESI MS/MS with a Thermo Scientific Easy-nLC 1000 (Thermo Scientific, Waltham, MA) coupled to a tribrid Orbitrap Fusion (Thermo Scientific, Waltham, MA) mass spectrometer. In-line de-salting was accomplished using a reversed-phase trap column (100 μm × 20 mm) packed with Magic C_18_AQ (5-μm 200Å resin; Michrom Bioresources, Auburn, CA) followed by peptide separations on a reversed-phase column (75 μm × 250 mm) packed with ReproSil-Pur 120 C_18_AQ (3-μm 120Å resin Dr. Maisch, Germany) directly mounted on the electrospray ion source. A 45-minute gradient from 2% to 35% acetonitrile in 0.1% formic acid at a flow rate of 300 nL/minute was used for chromatographic separations. A spray voltage of 2200 V was applied to the electrospray tip and the Orbitrap Fusion instrument was operated in the data-dependent mode, MS survey scans were in the Orbitrap (AGC target value 500,000, resolution 120,000, and injection time 50 ms) with a 3 sec cycle time and MS/MS spectra acquisition were detected in the linear ion trap (AGC target value of 10,000 and injection time 35 ms) using HCD activation with a normalized collision energy of 27%. Selected ions were dynamically excluded for 45 seconds after a repeat count of 1.

Data analysis was performed using Proteome Discoverer 2.5 (Thermo Scientific, San Jose, CA) using Sequest HT as the protein-database search algorithm. The data were searched against an Uniprot Mouse (UP000000589 from March 07, 2021) database that included common contaminants (cRAPome Jan 29, 2015). Searches were performed with settings for the proteolytic enzyme trypsin and maximum missed cleavages was set to 2. The precursor ion tolerance was set to 10 ppm and the fragment ion tolerance was set to 0.5 Da. Variable modifications included oxidation on methionine (+15.995 Da) and carbamidomethyl on cysteine (+57.021 Da). All search results were run through Percolator for peptide validation and peptide results were filtered to a 1% false discovery rate.

