## Supplemental Figures and Legends for "Oncogenic role of a developmentally regulated *NTRK2* splice variant"

**Pattwell_FigS1**

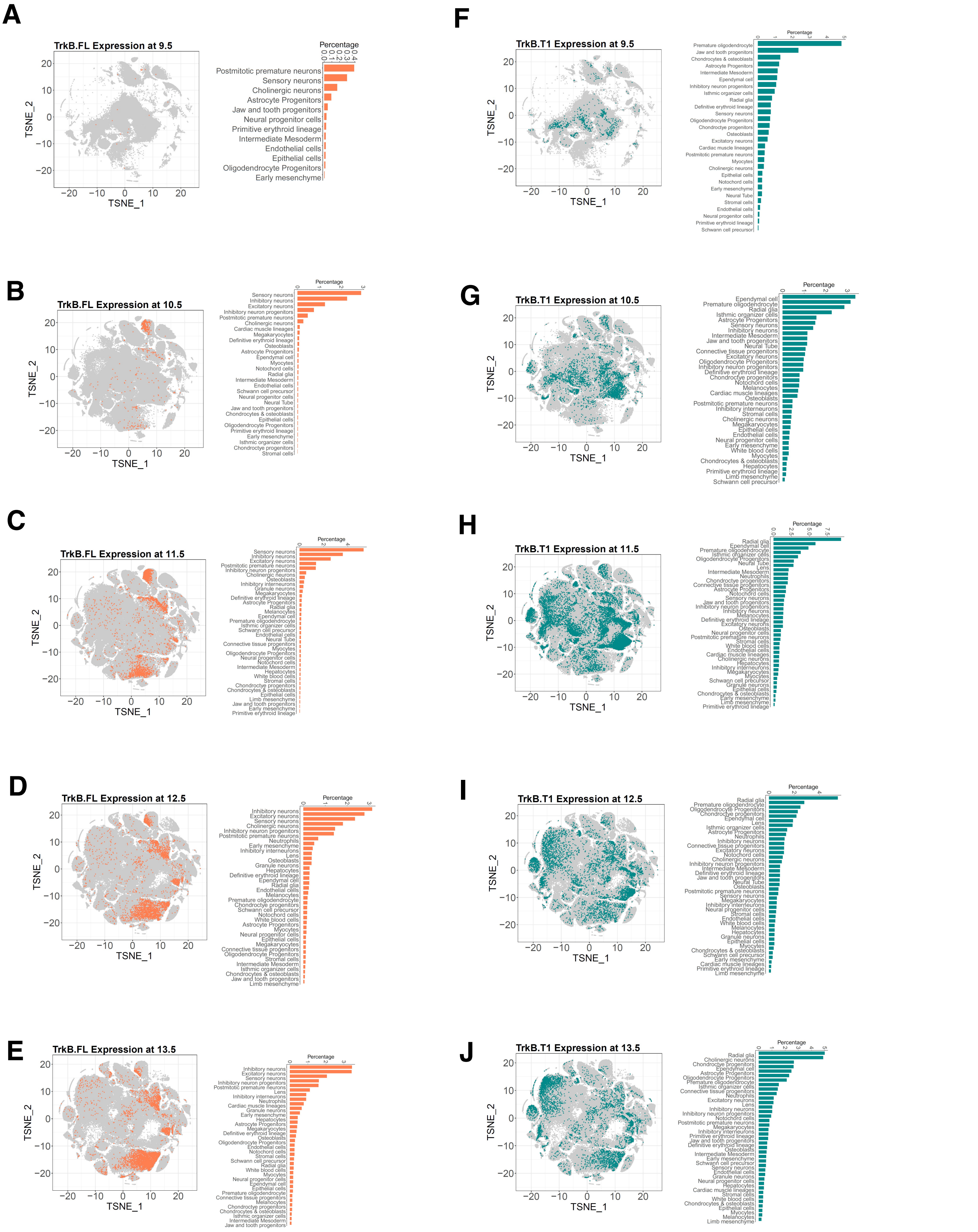


**Supplemental Figure S1: sci-RNA-Seq 3 transcript analyses across E9.5-E13.5 mouse embryonic development shows TrkB.FL expression in neuronal clusters with widespread TrkB.T1 expression in CNS and mesenchymal trajectories.** Visualization and percentages of cell types expressing TrkB.FL and TrkB.T1 across days: 9.5 **(A,F)**, 10.5 **(B, G)**, 11.5 **(C, H)**, 12.5 **(D, I)**, 13.5 **(E, J)**.

**Pattwell_FigS2**

A

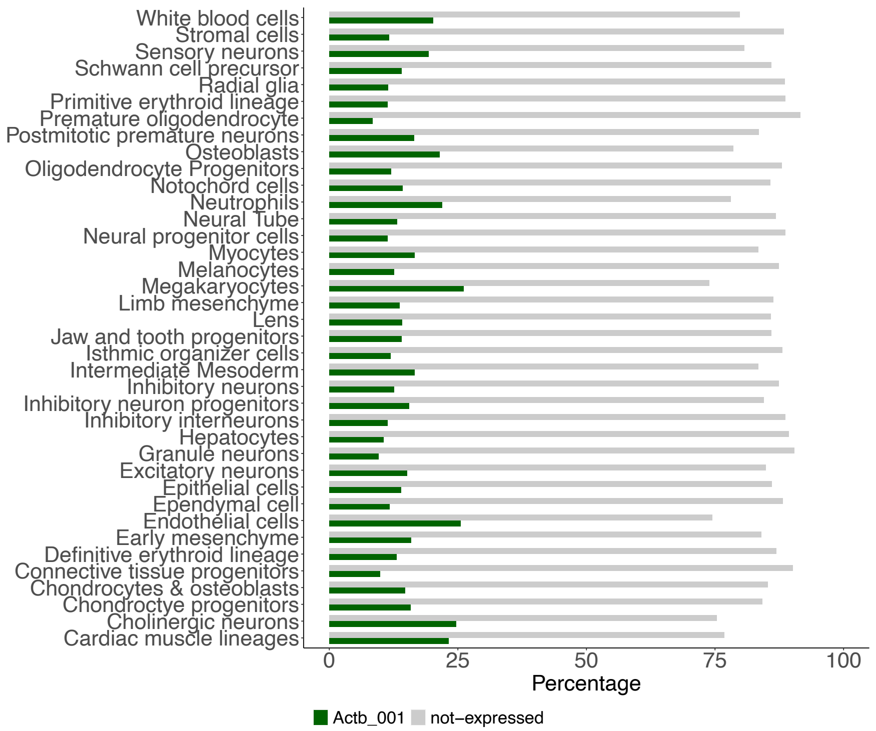


B

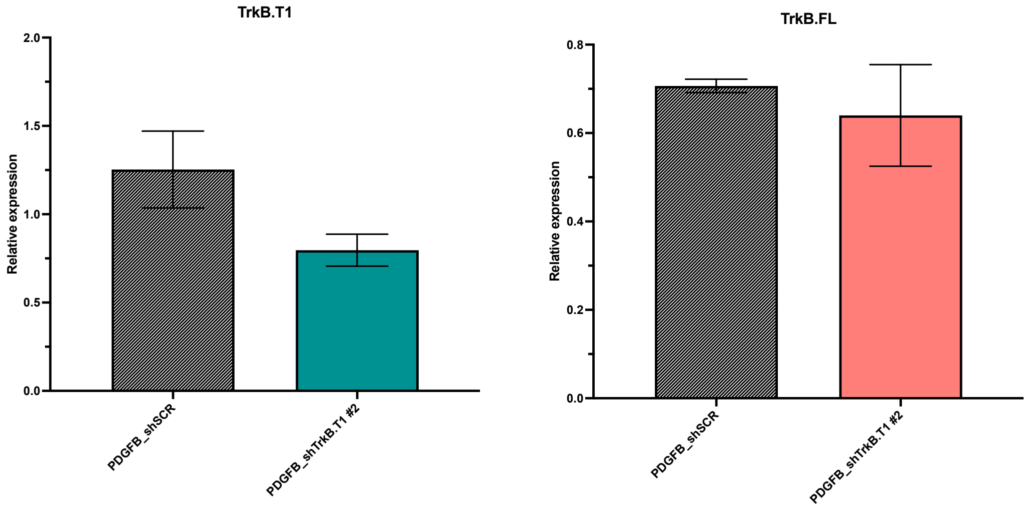


**Supplemental Figure S2: Control for sci-RNA-Seq3 and qRT-PCR. (A)** Quantification of beta-actin expression (*ACTB*) across the major MOCA cell types described by Cao et al. (2019). Beta-actin expression levels across all major clusters peak near 25% for cell types with the highest expression levels. **(B)** qRT-PCR for TrkB.T1 and TrkB.FL transcripts shows successful TrkB.T1 knockdown efficiency with no change in TrkB.FL levels for RCAS-PDGFB-shTrkB.T1 construct #2, which was used for subsequent analyses.

### Pattwell_FigS3

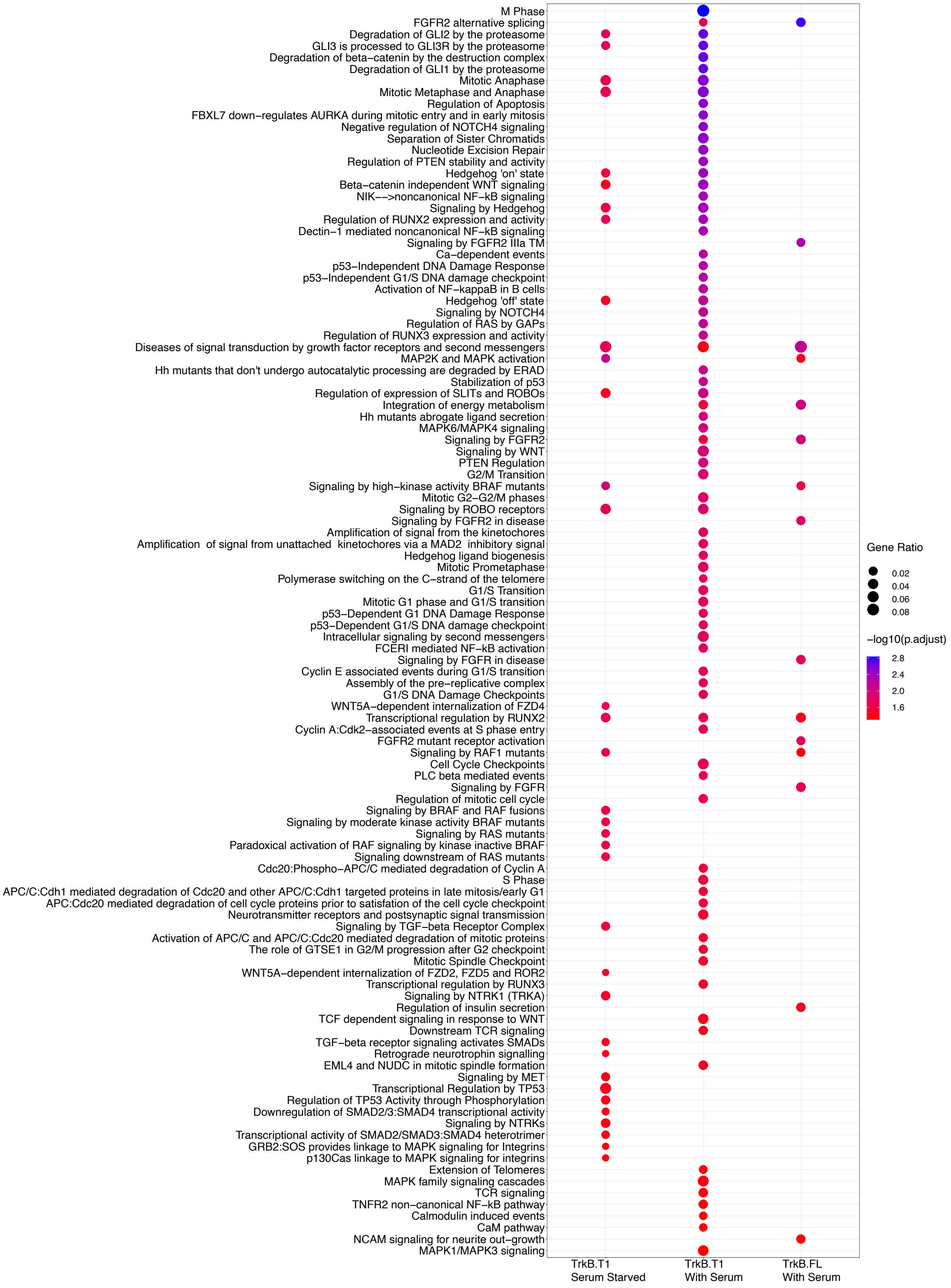


**Supplemental Figure S3: Proteomics analysis reveals developmental, oncogenic, and cell cycle/cell division pathways associated with TrkB.T1.** Gene ontology (GO) and Reactome Pathway analysis revealed distinct terms implicated in developmental, oncogenic, and cell cycle/cell division pathways under normal culture conditions (TrkB.T1 and TrkB.FL; with serum) and upon serum starvation (TrkB.T1 only; without serum).

### Pattwell_FigS4

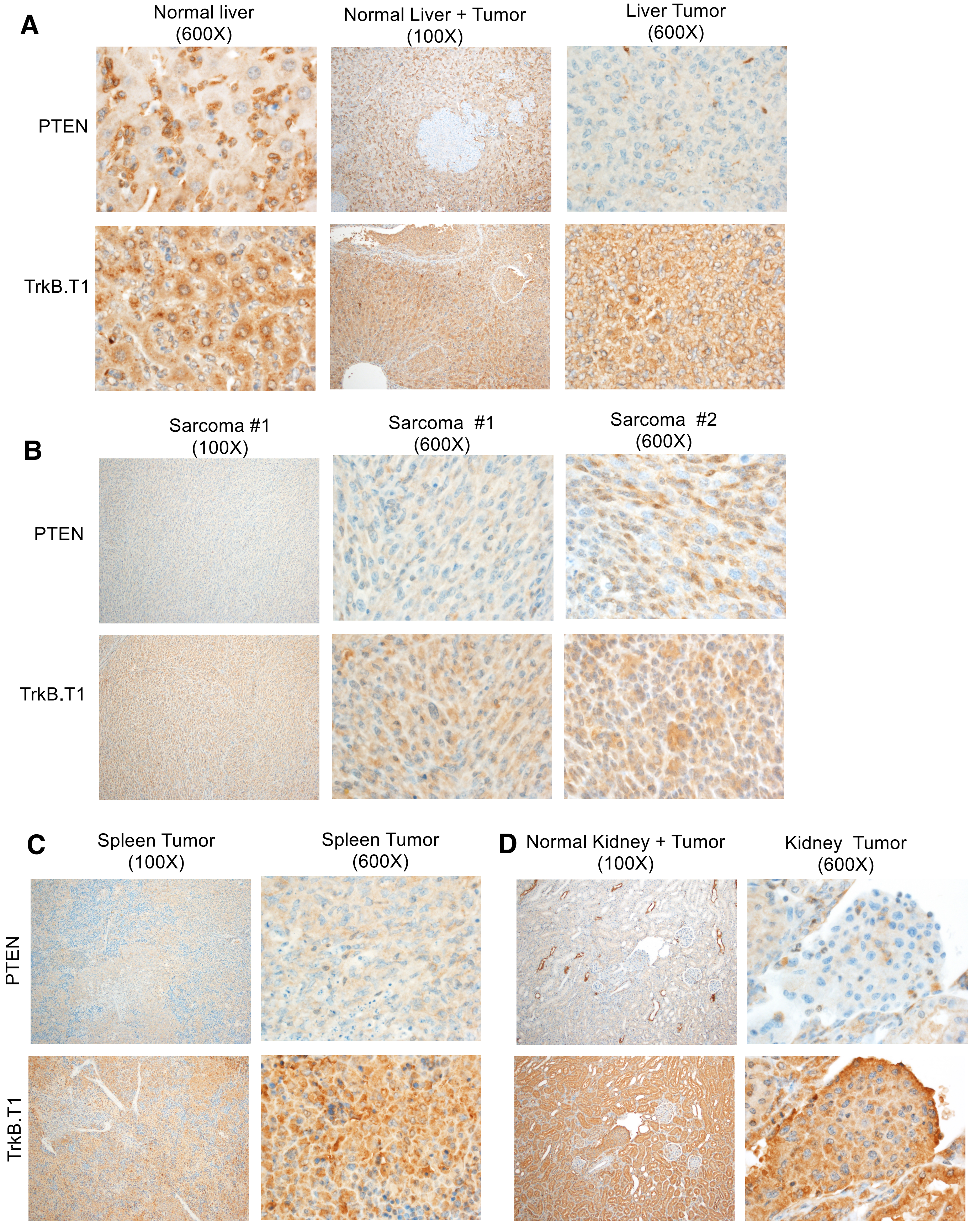


**Supplemental Figure S4: Representative images of PTEN and TrkB.T1 protein expression in mouse tumors.** Immunohistochemical staining reveals loss of PTEN and high expression of TrkB.T1 in tumors within the liver **(A)**, in sarcomas **(B)**, and in the spleen **(C)** and kidney **(D)**.

**Pattwell_FigS5**

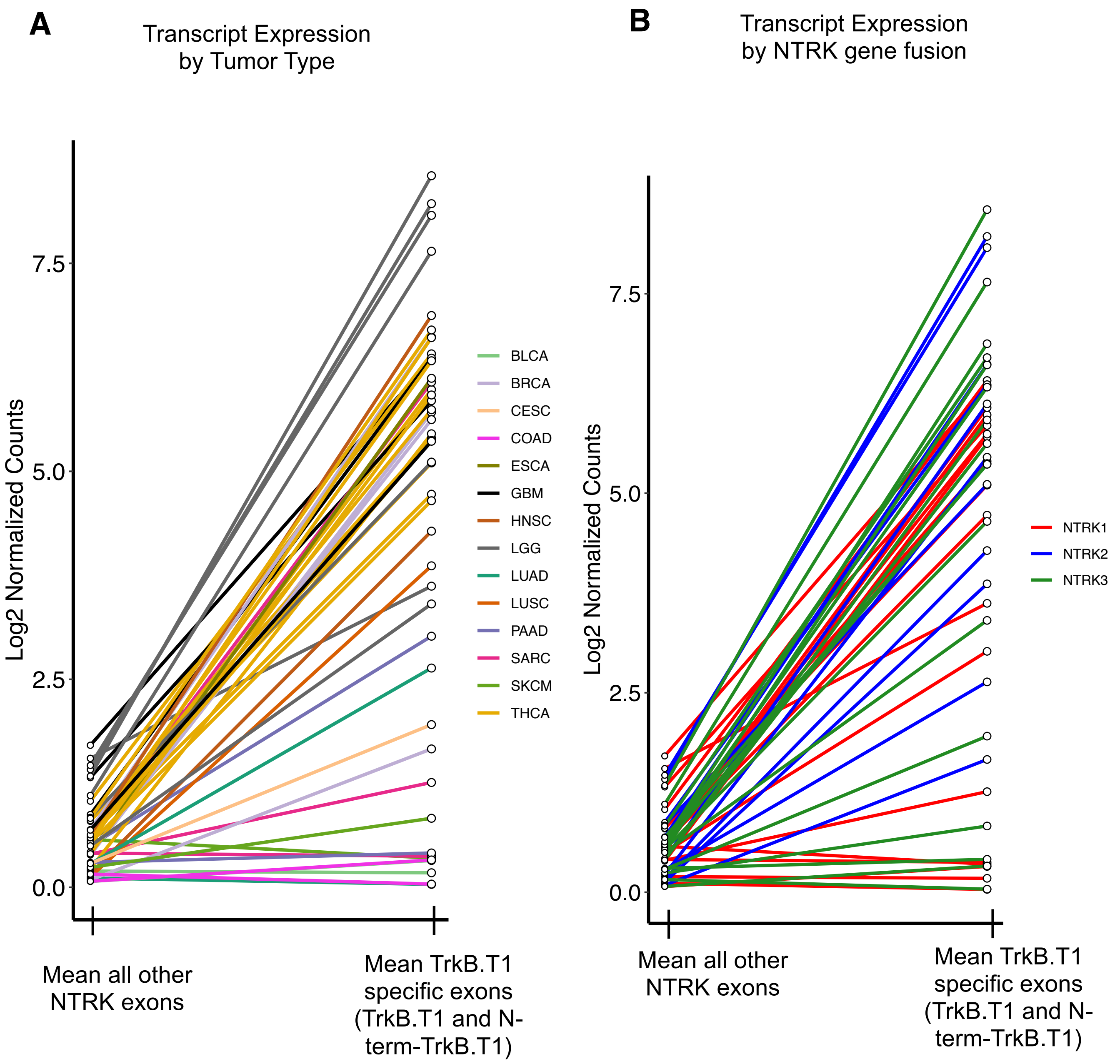


**Supplemental Figure S5: In depth transcript characterization of all TCGA tumors harboring known *NTRK* fusions**. Means are plotted for TrkB.T1-containing 11-amino acid specific exon (right) compared to means for all other *NTRK1, NTRK2, NTRK3* exons (left) characterized by tumor type (bladder urothelial cancer (BLCA), breast invasive carcinoma (BRCA), cervical squamous cell carcinoma and endocervical adenocarcinoma (CESC), colon adenocarcinoma (COAD), esophageal carcinoma (ESCA), glioblastoma multiforme (GBM), head & neck squamous carcinoma (HNSC), low grade glioma (LGG), lung adenocarcinoma (LUAD), lung squamous cell carcinoma (LUSC), pancreatic adenocarcinoma (PAAD), sarcoma (SARC), skin cutaneous melanoma (SKCM), thyroid carcinoma (THCA) **(A)** and *NTRK1, NTRK2,* or *NTRK3* fusion **(B)**.

#### Pattwell_FigS6

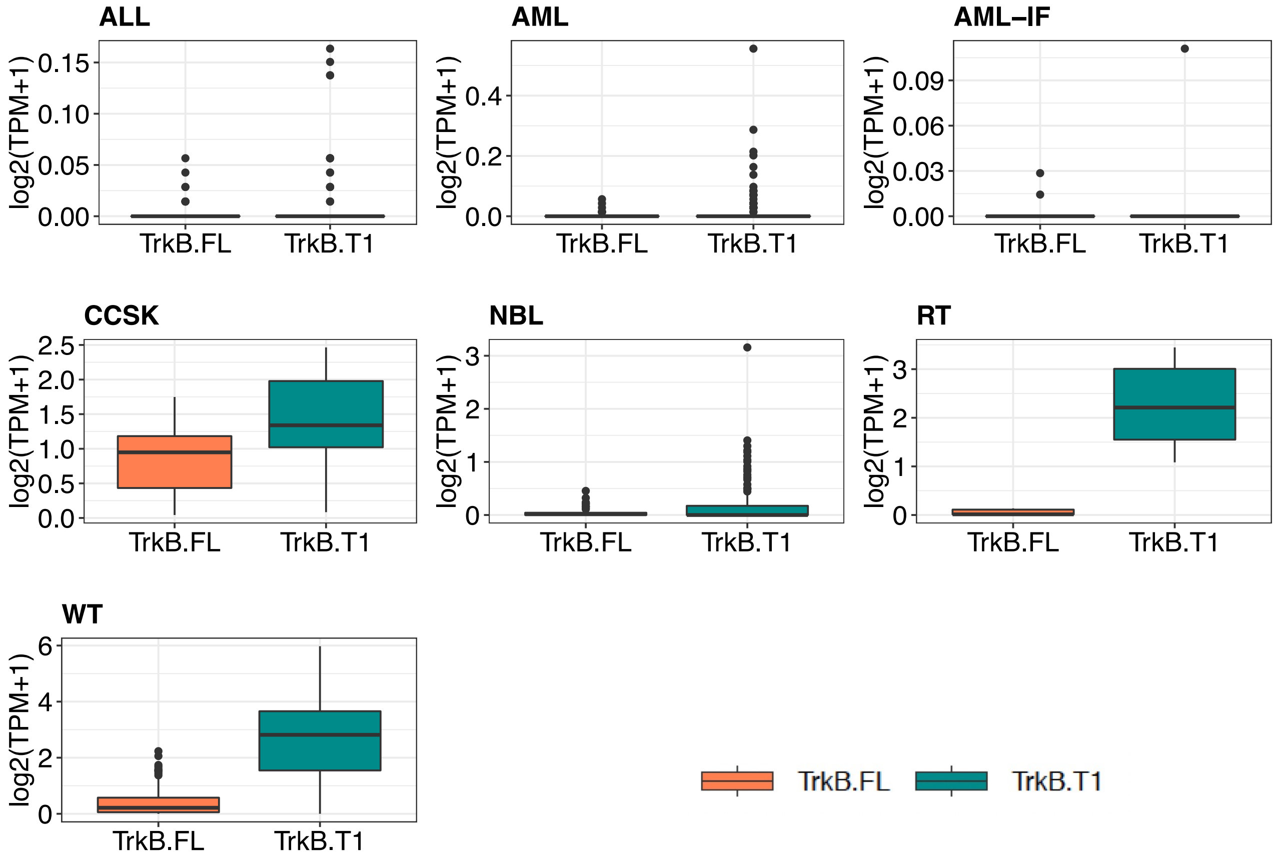


**Supplemental Figure S6: NTRK2 transcript analysis of pediatric data from the Therapeutically Applicable Research to Generate Effective Treatments (TARGET) program.** These data show that, similar to adult tumors in TCGA (as presented in **Fig. 5**) TrkB.T1 expression is the predominant isoform compared to TrkB.FL in several cancer types. Acute lymphocytic leukemia (ALL), acute myeloid leukemia (AML), induction failure AML (IF-AML), clear cell sarcoma of the kidney (CCSK), neuroblastoma (NBL), rhabdoid tumor (RT) and Wilms tumor (WT). Data are represented as boxplots where the middle line is the median, the lower and upper hinges correspond to the first and third quartiles (the 25th and 75th percentiles), the upper whisker extends from the hinge to the largest value no further than 1.5 * IQR from the hinge (where IQR is the inter-quartile range, or distance between the first and third quartiles) and the lower whisker extends from the hinge to the smallest value at most 1.5 * IQR of the hinge while data beyond the end of the whiskers are outlying points that are plotted individually
